# Intranasal Delivery of shRNA to Knockdown the 5HT-2A receptor Enhances Memory and Alleviates Anxiety

**DOI:** 10.1101/2023.12.27.573449

**Authors:** Troy T. Rohn, Dean Radin, Tracy Brandmeyer, Peter G. Seidler, Barry J. Linder, Tom Lytle, John L. Mee, Fabio Macciardi

## Abstract

Short-hairpin RNAs (shRNA) targeting knockdown of specific genes hold enormous promise for precision-based therapeutics to treat numerous neurodegenerative disorders. However, whether shRNA constructed molecules can modify neuronal circuits underlying certain behaviors has not been explored. We designed shRNA to knockdown the human *HTR2A* gene *in vitro* using iPSC-differentiated neurons. Multi-electrode array (MEA) results showed the knockdown of the 5HT-2A mRNA and receptor protein led to a decrease in spontaneous electrical activity. *In vivo*, intranasal delivery of AAV9 vectors containing shRNA resulted in a decrease in anxiety-like behavior in mice and a significant improvement in memory in both mice (104%) and rats (92%) compared to vehicle-treated animals. Our demonstration of a non-invasive shRNA delivery platform that can bypass the blood-brain barrier has broad implications for treating numerous neurological mental disorders. Specifically, targeting the *HTR2A* gene presents a novel therapeutic approach for treating chronic anxiety and age-related cognitive decline.

---

Neurological disorders such as Alzheimer’s disease (AD) and chronic anxiety are a major public mental health challenge, affecting millions of people worldwide. However, despite significant research efforts, there has been limited success in treating the symptoms associated with these disorders. Precision-based therapeutics such as CRISPR/Cas9 and RNA interference molecules offer a promising new approach to treating neurological and neurodegenerative disorders. shRNA represents one class of RNA interference molecules that has a mechanism based on the sequence-specific degradation of host mRNA through cytoplasmic delivery and degradation of double-stranded RNA through the RISC pathway ^1,2^. Whereas CRISPR/Cas9 leads to permanent changes in the genome, shRNA induces reversible gene silencing through a posttranslational regulatory process targeting degradation of specific mRNAs. Besides being reversible, shRNA has also been widely used in research for over a decade and there are currently four FDA-approved therapeutics that use shRNA to treat rare metabolic disorders ^3^. Moreover, shRNA targets a specific mRNA sequence, meaning it can potentially distinguish between closely related genes with high sequence homology. However, whether shRNA could be used to modify certain behavioral traits within the CNS has not been investigated.

Currently there is a great need for non-invasive methods of delivering gene therapy to the brain. shRNA represents a potential powerful tool, but it is difficult to deliver to the brain because of the blood-brain barrier (BBB). The BBB is a protective layer that prevents most molecules from entering the brain ^4^. One way to overcome this challenge is to use adeno-associated viral (AAV) vectors. AAV vectors are viruses that have been modified to be safe and effective for gene delivery due to their ability to deliver stable, long-lasting transgene expression in non-dividing cells ^5^.

We recently demonstrated that intranasal delivery of CRISPR/Cas9 encapsulated within adeno-associated viral (serotype AAV9) vectors could bypass the BBB and lead to knockout of the *HTR2A* gene in neuronal populations ^6^ (US Patent Application No. 63/283,150). The *HTR2A* gene encodes for the 5HT-2A receptor, one of the fifteen serotonin receptor subtypes expressed in the brain and is implicated in both anxiety disorders ^7, 8^ and memory ^9–11^. In the present study we designed a shRNA to knockdown the *HTR2A* gene and demonstrate a decrease in spontaneous electrical activity in human iPSC-differentiated neurons *in vitro* as well as enhanced memory and a reduction in anxiety in mice and rats *in vivo*. The development of this non-invasive shRNA delivery platform, which is capable of bypassing the blood-brain barrier, holds substantial implications for treatment of a wide spectrum of neurological and neurodegenerative disorders. Specifically, the targeting of the *HTR2A* gene emerges as a novel and promising therapeutic approach for addressing conditions such as chronic anxiety, mild cognitive impairment, dementia, and possibly AD.

## Materials and Methods

### Guide RNA and AAV9 vector design for CRISPR/Cas9 experiments

Details on the methods used to synthesize, design, and validate guide RNA (gRNA) and knockdown of the mouse *HTR2A* gene have been previously reported ^6^. In brief, the selected gRNA, TGCAATTAGGTGACGACTCGAGG (US Patent Application No. 63/283,150), would give no predicted off-target cut sites, produce an 86.6% frameshift frequency, and a precision score of 0.55. Two different adeno-associated virus serotype 9 (AAV9) vectors were designed to deliver spCas9 and gRNA to CNS neurons. The design of two vectors was necessary based on the limited carrying capacity of 4.7 kb for AA9 viruses ^12^. We created dual AAV9 systems to expand the capacity of an effective silencing of the target gene. The first AAV9 expressed spCas9 under a neuronal-specific promoter, MeCP2, and the spCas9 vector utilized the PX551 plasmid from Addgene (pAAV-pMecp2-SpCas9-spA) ^13^. The second AAV9 vector consisted of the gRNA sequence and a green-fluorescence protein (GFP) reporter under the U6 promoter (AAV-GFP-ssODN-U6-gRNA) ^6^. Titer load (in genome copy number per ml, or GC/ml) was determined through quantitative real-time PCR, with typical yields giving 2.0 x 10^13^ GC/ml. Both AAV9 vectors were stored in phosphate buffered saline (PBS) with 5% glycerol at -80°C until used. Design, manufacturing, and purification of the AAV9 vectors used in this study were performed by Vector Biolabs (Malvern, PA).

### shRNA design and AAV9 vector design

Different strategies were employed depending on whether CRISPR/Cas9 or shRNA was utilized. For the CRISPR/Cas9 experiment, we proceeded as described in the previous paragraph. For CNS delivery of shRNA, a similar approach was undertaken, but in this case a single AAV9 vector was used. It consisted of a single DNA plasmid containing the following target shRNA sequence to the human HTR2A RNA: (US Provisional Patent Application Serial No. 63/470,150):

GCTGTTCTGAAGACAAAGAACTCTGGTTTTGGCCACTGACTGACCAGAGTTCTGTCTTC AGAA CAG The human *HTR2A* gene consists of four exons that give rise to two major isoforms and is found on chromosome 13. The predicted binding region of the primary RNA transcript for this sequence is the beginning of exon 4, which would lead to the potential knockdown of all possible isoforms (Fig. 1A). This specific shRNA sequence was chosen based upon validation and screening of four different shRNA sequences. As shown in Fig. 1B, in contrast to the empty vector control, Shmir#3 led to an 87% knockdown of the targeted RNA sequence. This sequence included three elements necessary for the construction of the complete shRNA molecule (**Supplementary Information, Figure 1**): 1) The targeting sequence TCTGAAGACAAAGAACTCTG; 2) The stem-loop feature of the shRNA; 3) The passenger strand. The RISC complex initially recognizes a double-stranded short interfering RNA, but only one strand is finally retained in the functional ribonucleoprotein complex. The non-incorporated strand, or ‘passenger’ strand, is removed during the assembly process and most probably degraded thereafter ^14^. In addition to the human shRNA construct, a scrambled control, AAV9-MeCP2-GFP-scrmb-shRNA, containing the identical target sequence but in random order, was synthesized in an identical manner.

**Fig. 1.**
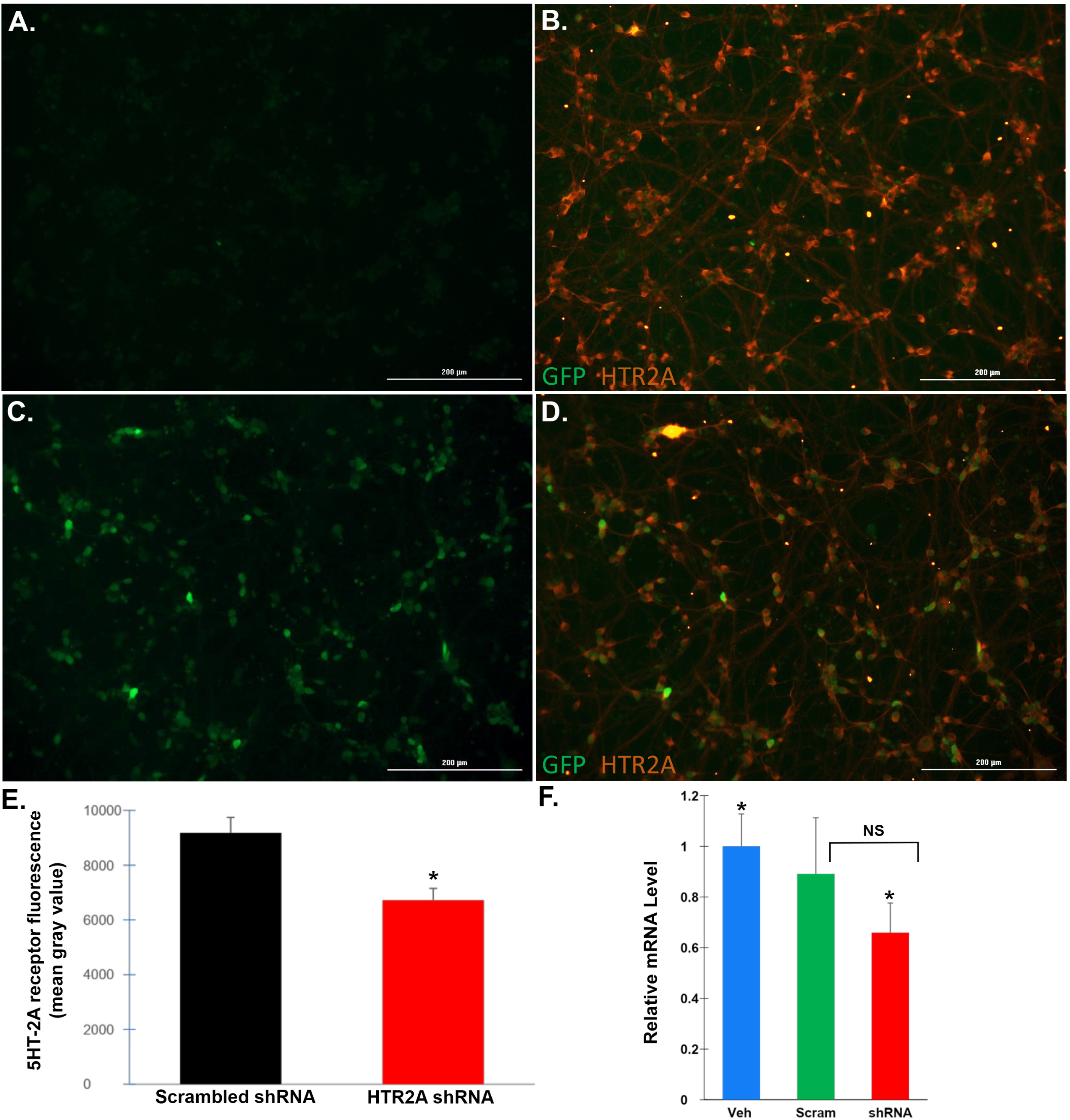
Treatment of human iPSC-differentiated glutamatergic neurons with shRNA leads to downregulation of the 5HT-2A receptor protein. **(A-D)**: Representative immunofluorescence images in human neurons following a 10-day treatment with scrambled AAV9 shRNA-AAV9 viral particles (A and B) or hu-HTR2A shRNA-AAV9 viral particles at MOI of 3×10^5^ **(C and D)**. Green fluorescence represents green fluorescence protein expression, while red fluorescence is indicative of 5HT-2A receptor protein following immunocytochemistry using an anti-mouse 5HT-2A receptor antibody at 1:50. Low level GFP expression, which served as a proxy for scrambled shRNA expression was observed in all cases (Fig. 1A). Panel B indicates robust expression of the 5HT-2A receptor protein in both cell bodies and neurites following treatment with the scrambled control. Panels C and D are representative images following treatment with hu-HTR2A shRNA-AAV9 and revealed stronger GFP expression **(C)**. **(D)**: Following hu-HTR2A shRNA-AAV9 treatment, a decrease in 5HT-2A fluorescent intensity was apparent. **(E)**: ImageJ quantification of 5HT-2A immunofluorescence indicated a significant 27% decrease in hu-HTR2A shRNA-AAV9-treated neurons as compared to scrambled-treated (*p-value = .0019, N= 3 different samples for each condition). **(F)**: Data show the results of qPCR real-time assays to analyze mRNA levels of Htr2a following extraction of RNA iPSC differentiated neurons in either vehicle-controls (blue bar) or scrambled treated (green bar), and hu-HTR2A shRNA (red bar). Neurons were treated on day 1 at a cell density of 200K/well and cells pelleted and frozen on day 16. Results display the relative change in expression using GAPDH as an internal control. Real-time PCR results represent a total of 3 separate treatments for each condition in which cells were pooled and frozen at -80°C. PCR experiments were performed in triplicate. The results indicated a significant 34% decrease in htr2a mRNA expression as compared to vehicle controls, where *denotes significant difference between the two groups, p = .012. Relative mRNA levels were not significantly different between vehicle control and scrambled-treated cells (p-value = .502) or between scrambled- and shRNA-treated neurons (p-value = .094).

For construction of the mouse shRNA to target knock-down of the 5HT-2A receptor, a similar approach was utilized. The mouse *HTR2A* gene encodes a single protein-coding transcript, Htr2A-201 located on chromosome 14 (**Supplementary Information, Figure 1**). The following sequence was used for assembly of the shRNA based on *in vitro* testing indicating a 77% knockdown:

> GCTGAGCACATCCAGGTAAATCCAGGTTTTGGCCACGACTGACCTGGATTTCTGGATG TGCT CAG

No knockdown was observed with the empty vector control or a scrambled shRNAmir control (**Supplementary Information, Figure 1**). For validation and screening, knockdown was verified using HEK293 cells co-transfected with the cDNA plasmid containing the *HTR2A* gene target. It is noteworthy that the designed shRNA target sequence for mice is 100% conserved in the rat *HTR2A* gene, thus this same construct was used in our rat studies.

For all designed shRNA delivery subcloning of the shRNA was carried out in a modified pAAV cis-plasmid under the neuronal-specific promoter, MeCP2. A reporter gene enhanced green fluorescent protein (eGFP) was subcloned upstream of the shRNA sequence. AAV9 viral large-scale transfection of plasmids was carried out in HEK293 cells and purified through a series of CsCl centrifugations. Titer load (in genome copy number per ml, or GC/ml) was determined through quantitative real-time PCR, with typical yields giving 1-2 x 10^13^ GC/ml. All AAV9 vectors were stored in PBS with 5% glycerol at -80°C until used. Design, manufacturing, and purification of AAV9 vectors used in this study were performed by Vector Biolabs (Malvern, PA).

### Culturing of human iPSC differentiated neurons

Human iPSC differentiated cortical glutamatergic neurons were cultured for immunocytochemistry (ICC), PCR analysis, and functional measurements using a multielectrode array (MEA) assay. Seeding density on 24-well Axion MEA plates was carried out at 90K/well to obtain confluent monocultures. Post thaw viability was 91.6%. Cells were thawed in seeding medium (**Supplementary Information, Table 1**) and then plated in 10 µl droplets at 90K/well, incubated at 37 °C for 20 minutes, and then 490 µL of seeding media was gently added to each well. On day 4 *in vitro*, a 500 µL media addition was performed with Day 4 Medium (**Supplementary Information, Table 2**). From day 7 to day 28 a 50% media change was performed every 3-4 days with Maintenance Medium (**Supplementary Information, Table 3**).

### Immunocytochemistry of human iPSC differentiated cortical glutamatergic neurons

A select group of cortical glutamatergic neurons were cultured and treated with scrambled or hu-HTR2A shRNA test articles (MOI 1.0×10^4^-1.0×10^6^) for an endpoint evaluation using ICC and GFP fluorescence to determine an optimal multiplicity of infection (MOI). Neurons were cultured in a 96-well plate format for a minimum of 10 days and the GFP signal was observed to determine optimal MOI of hu-HTR2A shRNA. Additionally, immunocytochemistry targeting the 5HT-2A receptor was performed and images were reviewed to determine the appropriate antibodies and dilutions. Based on these preliminary studies, we chose the human specific 5HT-2A antibody from ThermoFisher PA5-120747 at 1:50 with secondary Goat Anti-Rabbit Fluor555 at 1:1000 using a MOI of 3 ×10^5^. For ICC, at 10 days *in vitro*, immunocytochemistry was initiated to stain for the 5HT-2A receptor. Wells were washed with 100 µL of 1X DPBS (PBS) for 5 minutes followed by fixation in 50 µL of 4% Paraformaldehyde (PFA) for 10 minutes. Once fixed, wells were rinsed twice with PBS for 5 minutes each and then blocked in 50 µL of 2.5% Donkey Serum (DS)/0.1% Triton in PBS for 10 minutes. After the blocking step was completed, 50 µL of blocking solution containing 5HT-2A receptor primary antibodies in PBS containing 2.5% DS and 0.1% Triton was added and left at 4°C overnight. Following overnight incubation in primary antibody, the primary antibody solution was removed, and target wells were rinsed twice with PBS. Next, a 5% DS in PBS was added to wells for 10 minutes. Following primary removal the secondary antibody solution containing Fluor 647 anti-chicken, Fluor 555 anti-rabbit, and Hoechst 40045 in a 5% DS in PBS was added to each well. Wells containing the secondary antibody solution were incubated at ambient temperature for 45 minutes protected from light. Once incubation was complete, wells were washed three times with PBS and imaging began. Additional fixed wells were stained with antibodies within 1 week to acquire additional reference images. Plates were stored at 4°C. Imaging was completed using a BioTek® LionHeart FX using 4X, 10X, or 20X objectives and excitation at 365nm, 465nm, 523nm, and 590nm. Phase contrast imaging was also completed using the same imaging system. Regions of interest (ROI) were identified within each MOI group to select regions showing clear neuron morphology and representative images were acquired.

### Functional assay using multi-electrode array (MEA) on monocultures of human iPSC differentiated glutamatergic neurons

Cells were thawed in seeding medium and plated at 90K/well on a CytoView MEA 24-White Plate as described above. Three different conditions were tested in quadruplicate starting at Day 0 (first day of plating): 1) Vehicle control consisting of PBS only; 2) Scrambled AAV9 shRNA at MOI of 3 x 10^5^; 3) hu-HTR2A shRNA at MOI 3 x 10^5^. Electrophysiological recordings were acquired three times per week following day 5 *in vitro* on the Axion Maestro Edge Platform. Each plate consisted of 24-wells with 16 electrodes in a 4×4 grid/well for a 384-channel configuration. The Maestro Edge was equilibrated to 37 °C and 5% CO2 prior to an MEA plate being placed on the instrument. Each plate was then equilibrated for 15 minutes, after which a 15-minute recording was taken from plate 92-0232 from Day 3 until day 30. The Day 14 recording showed an overwhelming anomaly in the signal, in every well on every electrode, indicating interference. Due to the interference, the data from this recording was excluded from analysis. Raw data files (*.raw) and spike files (*.spk) were recorded using the software AxIS Navigator (s 1.5.1.17) on the Spontaneous Neural Configuration setting. Neuronal spikes were detected using an adaptive thresholding set to 6 times standard deviation (6SD) of the mean noise level. Each *.spk was loaded into Neural Metric Tool (Version 2.4.12, Axion Biosystems) for data analysis (*.csv) and to obtain spike raster plots. Active electrodes (AEs) were defined as >1 spikes/min. Bursting electrodes were defined using inner spike interval (ISI) parameters set to a minimum number of spikes of 5 per ISI event and maximum ISI of 100 milliseconds (ms). Network bursting was extracted using the minimum of 50 spikes and 35% of electrodes.

Each graph, mean firing rate (MFR), electrode and network Bursting, and Synchrony Index, was generated using data from *.csv files in the software Origin (Pro), version 2022. All culturing and characterization of iPSC neurons as well as immunohistochemistry and MEA analysis were performed by BrainXell (Madison, WI, USA).

### Quantitative real-time qPCR

A select group of human iPSC differentiated cortical glutamatergic neurons were cultured and treated with scrambled AAV9, hu-HTR2A shRNA AAV9, or D-PBS vehicle with an endpoint of collecting cell pellets. The neurons were cultured in a 24-well plate, with AAV added to the wells within one hour of seeding at a density 200K/well. Cells were cultured to 16 days *in vitro*, at which time they were dissociated from the wells and pelleted via centrifugation. Pelleted cells were immediately frozen at -80C. Total RNA was extracted from frozen cells using a standard extraction protocol with Trizol, dissolved in DEPC-treated deionized water and quantified. Following reverse transcription, qPCR was carried out using the following primers:

**Primer Sequence (5’->3’)** 1. **GAPDH** Forward: TGAAGGTCGGAGTCAACGGAT Reverse: CCTGGAAGATGGTGATGGGAT **2. HTR2A** Forward: CTTCCAGCGGTCGATCCATAG Reverse: GCAGGACTCTTTGCAGATGAC

The relative expression was determined by calculating the 2^-Δct^ value. The 2^(-ddCt) value was calculated and normalized to calculate a fold difference between the vehicle controls and the AVV9-treated groups. The RNA extraction and qPCR were performed by Creative Biogene (Shirley, NY, USA).

### Test formulation and intranasal administration for *in vivo* delivery

For CRISPR/Cas9 T-maze alternation task behavioral studies, the AAV-treated group consisted of an equal mixture of AAV9-gRNA-U6-GFP and AAV9-Mecp2-spCas9 suspended in 0.9% NaCl (saline). The concentration of AAV9 stock mixture was ∼5.0 ×10^12^ GC/ml for mice tested. For each behavioral test, a new, independent batch of AAV9 vectors was synthesized and prepared accordingly. The AAV cocktail was administered twice on day one (morning and afternoon) 5 weeks before the execution of the behavioral tests. For intranasal delivery of AVV, mice were hand-restrained with the nose positioned to facilitate the dosing. A meniscus of AAV solution droplet (10 µl per nostril) was then formed at the tip of the micropipette and presented for inhalation in each of the nares of the mouse. Each mouse (N=15) received a total of 40 µl of AAVs equivalent to ∼2 x 10^11^ viral particles, whereas vehicle-treated animals (N=15) received 40 µL of saline for each treatment following the same protocol.

To test shRNA targeted to knockdown the *HTR2A* gene *in vivo*, 100 µL of AAV9-MeCP2-GFP-mHTR2A-shRNAmir at 1-2×10^13^ GC/ML was mixed with 100 µL of saline. In that solution the stock AAV9 was at ∼6.5×10^12^ GC/ML. Animals (N= 15 for each group) were then dosed with 40 µl AAV9-MeCP2-GFP-mHTR2A-shRNA as a single 20 µL dose (10 µl per nostril) twice on day 1, five weeks before the first trial. Thus, each mouse received a total of 40 µl of AAVs equivalent to 1-2 x 10^11^ viral particles. Vehicle treated animals received 40 µl of saline for each treatment. For both groups, mice were treated on day 1 and assessed behaviorally 5 weeks later, unless otherwise specified.

### Light dark behavioral test

The light dark test was performed at one timepoint on the 5th and 8th week after the treatment using 2-month-old CD-1 male mice. In this task, mice were given a choice between exploring a brightly lit chamber or a dark chamber as a measure of anxiety. The apparatus consisted of two PVC (polyvinylchloride) boxes (19 × 19 × 15 cm) covered with Plexiglas. One of these boxes was darkened. The other box was illuminated by a desk lamp placed above and providing an illumination of approximately 2000 Lux. An opaque plastic tunnel (5 × 7 × 10 cm) separated the dark box from the illuminated one. A camera linked to a video tracking system (Viewpoint, France) was used to monitor the behavior of the mouse in the lit box. Animals were placed individually in the lit box, with their heads directed towards the tunnel. The time spent in the lit box and the number of transitions between the two boxes was recorded over a 5 min period after the first entry of the animal in the dark box. The total walked distance in the lit box was also recorded. The apparatus was cleaned between each animal using 70% alcohol. A total of 15 mice were used for each group.

### T-maze continuous alternation task (T-CAT)

The T-maze continuous alternation task (T-CAT) is among the methods implemented to evaluate the spatial exploratory performance in rats or mice ^15^. It relies on spatial and working memory and is sensitive to various pharmacological manipulations affecting memory processes. Exploratory studies performed at Neurofit SAS (Illkrich, France) indicate that cognitive dysfunction occurs in aged male C57Bl6 mice (12 months old) and that this deficit can be reversed by drugs with cognitive enhancing properties such as nicotine and donepezil. The aim of this study was to investigate the potential cognitive enhancing properties of 5HT-2A receptor knockdown using CRISPR/Cas9 on aged (12 months old) mice with an age-dependent cognitive dysfunction. Aged male C57Bl6 mice (12 months old) were used and randomly distributed to control or experimental groups (15 animals per group). The T-maze consisted of 2 choice arms and 1 start arm mounted to a square center. Sliding doors were provided to close specific arms during the forced choice alternation task. During the trials, animal handling and the visibility of the operator were minimized as much as possible. The experimental protocol for this task consists of a single session which starts with one “forced-choice” trial, followed by 14 “free-choice” trials. In the first “forced-choice” trial, the animal is confined 5 s in the start arm and then it is released while either the left or right goal arm is blocked by closing the sliding door. Afterwards, mice negotiate the maze at will, eventually entering the open goal arm and returning to the start position. Immediately after the return of the animal to the start position, the left or right goal door is opened, and the animal is allowed to choose freely between the left and right goal arm (“free choice” trials). The animal is entered in an arm when it places its four paws in the arm. A session is terminated, and the animal is removed from the maze as soon as 14 free-choice trials have been performed or 15 min have elapsed, whichever occurs first. The apparatus is then cleaned between each animal using 70% alcohol. The percent of spontaneous alternations between the two arms is calculated as the number of spontaneous alternations divided by the number of free-choice trials.

### Novel object recognition test

The object recognition task is used to assess short-term memory, intermediate-term memory, and long-term memory in rats ^16^. The task is based on the natural tendency of rats to preferentially explore a novel versus a familiar object, which requires memory of the familiar object. The time delay design allows for the screening of compounds with potential cognitive enhancing properties to overcome the natural forgetting process. To test whether shRNA-knockdown of the rat 5HT-2A receptor improved memory, Wistar male rats (12 animals per group) were randomly assigned to two groups consisting of vehicle (PBS) or AAV9-MeCP2-GFP-mHTR2A shRNA. Following administration of the vehicle or AAV9 compound (see above), animals were assessed in this task at both 3 and 5 weeks later.

### The behavioral protocol consists of 4 steps

Step 1 - Habituation: 24 hours before the first trial, animals are habituated to the apparatus for 15 min.
Step 2 -Acquisition: Object A is placed at the periphery of a central square (∼30 × 30 cm). Memory acquisition session lasts for 10 minutes.
Step 3 -Retention: 24 hours later, objects A (familiar) and B (novel) are placed at two adjacent locations of the central square. The number of contacts and time spent in contact with the objects are recorded.
Step 4 -Recognition: For each animal, the time taken to explore object A (t_A_) and object B (t_B_) are used to create a recognition index (RI) determined as RI = t_B_/(t_A_ + t_B_) x 100.

The arena and objects were cleaned with 70% alcohol between each rat test session. These behavioral studies were performed by Neurofit SAS. All animal care and experimental procedures were performed in accordance with institutional guidelines and were conducted in compliance with French Animal Health Regulation.

For all behavioral studies, animals were keyed, and data were blinded until the end of experiments.

### Tissue preparation

Immediately following behavioral analysis, mice or rats were anesthetized with 5% isoflurane/oxygen mixture and sacrificed by decapitation. Brains, including the olfactory bulbs, were extracted, and fixed in 4% formalin for 48 hours and transferred to vials containing 1% formalin in PBS buffer. Brain samples were stored at 4°C. Alternatively, brains were flash frozen and stored at -80°C for RNA extraction and analysis by PCR.

### Statistical analysis

Behavioral data were analyzed by independent sample t-tests using JASP (Version 0.17.3, University of Amsterdam), and microelectrode array data were analyzed via repeated measures ANOVAs using Statistica (Version 13.5, Tibco Software). All data used in these tests were checked and found to conform to parametric assumptions.

### Quantitative real-time qPCR in mice or rat brain tissue

Total mouse brain RNA was extracted from frozen brains using a standard extraction protocol with Trizol, dissolved in DEPC-treated deionized water and quantified. Following reverse transcription, qPCR was carried out using the following primers:

Prime-F: 5’-AGAGGAGCCACACAGGTCTC-3’ and

Primer-R: 5’-ACGACAGTTGTCAATAAAGCAG-3’. The relative expression was determined by calculating the 2^-Δct^ value. RNA extraction and qPCR was contracted out to Creative Biogene (Shirley, NY, USA).

The following primers were used:

> Rat Htr2a-F: 5’-CACCGACATGCCTCTCCAT-3’

> Rat Htr2a-R: 5’-AGGCCACCGGTACCCATAC-3’

> Rat-GAPDH-F: 5’-TGGCCTCCAAGGAGTAAGAAAC-3’

> Rat-GAPDH-R: 5’-GGCCTCTCTCTTGCTCTCAGTATC-3’

### RNA Isolation Method

Tissue samples were pulverized using a sterilized mortar and pestle in the presence of liquid nitrogen. Samples were then transferred to a new, chilled tube and 1 mL of TRIzol reagent was added and thoroughly mixed. After the incubation period, 0.2 mL of chloroform was introduced to each tube. Subsequently, the samples were centrifuged at 12,000 x g for 15 minutes at a temperature of 4°C. RNA was transferred to a new tube and 0.5 mL of isopropanol was added to this aqueous phase and incubated for 10 minutes at 4°C. RNA pellets were then resuspended in 1 mL of 75% ethanol. After brief vortexing, the samples were centrifuged again at 7500 x g for 5 minutes at 4°C, and the supernatant was removed with a micropipette. The RNA pellet air dried for a period of 5 to 10 minutes. Finally, the dried RNA pellet was resuspended in a volume of 20 to 50 µL of RNase-free water.

### qPCR Method

For the qPCR process, 2 µg of total RNA was utilized, and cDNA was synthesized according to the kit manufacturer’s recommendations. This cDNA solution was then diluted by adding a 9-fold volume of water. For setting up the qPCR reaction, 17 µL of the master mix was dispensed into each well of a 96-well plate using a multichannel pipette. Separately, 3 µL of cDNA was added to each well. Each well, therefore, contained 5 µL of cDNA from different samples, 0.2 µL of a qPCR Primer Mix at 10 µM concentration, 10 µL of SYBR GREEN Mix at 2x concentration, and 4.8 µL of water, bringing the total volume to 20 µL. Samples were run at 95°C for 5 minutes, followed by 40 cycles of 95°C for 15 seconds and 60°C for 60 seconds.

### Immunohistochemical fluorescence microscopy

Following dehydration, 4 µm paraffin-embedded, sagittal sections were cut just lateral to the midline and used for immunofluorescence labeling. Briefly, all tissue sections were labeled with anti-GFP antibody (rabbit mAB #2956) 1:1,000 (Cell Signaling Technology, Inc., Danvers, MA, USA) or anti-5HT-2A receptor antibody (rabbit polyclonal, #24288) at 1:500 dilution (Immunostar, Hudson, WI). Secondary antibodies were conjugated to FITC or Cy3. DAPI was used as a nuclear stain. Whole slide scanning was performed using a Pannoramic Midi II scanner using a 40X objective lens with optical magnification of 98X, 0.1 µm/pixel. All sectioning, immunolabeling, and capturing of images was contracted out to iHisto (Salem, MA).

### ImageJ quantification

The level of 5HT2A receptor fluorescence was quantified using ImageJ software. This was accomplished by capturing 2X immunofluorescence images from three separate tissue sections from each group (vehicle control or AAV-treated). All data were expressed as the mean gray value ±SEM. The mean gray value reflects the sum of the gray values of all the pixels in the selected area divided by the number of pixels. The area of selection in square pixels was identical between vehicle-controls and AAV-treated for all analyses.

## Results

### Delivery platform of htr2a-shRNA utilizing adeno-associated viruses

*In vitro* analysis of designed shRNAs indicated efficient knockdown of the mouse/rat (77%) and human form (87%) mRNA. Following validation of both shRNAs, we developed an *in vivo* delivery platform for these reagents. Adeno-associated viruses (AAVs) are a popular choice for delivering shRNA cargo due to their low immunogenicity, good safety profile, and ability to achieve long-term expression in non-dividing cells ^12^. We selected the AAV9 serotype, which has been shown to be a highly efficient vector for transgene expression in neurons throughout the CNS ^17, 18^. A single AAV9 vector has the storage capacity (4.7 kb) to hold the DNA plasmid containing either the mouse, rat, or human htr2a shRNA. Since mouse and rat show 100% identity of the target sequence within the *HTR2A* gene, the same DNA plasmid construct was used for either species. Both constructs (mouse/rat and human) contained the DNA sequencing necessary for full assembly of a shRNA molecule following infection and delivery within neurons. To ensure neuronal specificity, expression of shRNAs was under the control of the neuronal specific promoter, MeCP2. AAV9 vector constructs also contained the GFP receptor gene to provide a visual proxy of shRNA expression. Typical viral titers were on the order of 1-2 x 10^13^ GC/ml. Herein, the mouse/rat and human shRNAs will be referred to as AAV9-MeCP2-GFP-mHTR2A-shRNA and AAV9-MeCP2-GFP-huHTR2A-shRNA, respectively.

### Adeno-associated virus exposure of human iPSC-differentiated glutamatergic cortical neurons with AAV9-MeCP2-GFP-huHTR2A-shRNA leads to a decrease in 5HT-2A receptor expression

As an initial approach, we tested *in vitro* the ability of AAV9-MeCP2-GFP-huHTR2A-shRNA to knockdown the human 5HT-2A receptor. Preliminary experiments were carried out with AAV9-MeCP2-GFP-huHTR2A-shRNA at various concentrations of viral particles to number of neurons (e.g., MOI). Results indicated an optimal MOI of 3 x 10^5^ on Day 1 *in vitro* and subsequent 5HT-2A receptor density was examined by immunocytochemistry following fixation on Day 10 using a human anti-5HT2A receptor antibody. As a control, we also tested a scrambled version of the targeted sequence packaged in an identical DNA plasmid within AAV9 vectors. As shown in Fig. 1A-D, treatment of neurons with AAV9-MeCP2-GFP-huHTR2A-shRNA led to a decrease in the expression of the 5HT-2A receptor as compared to scrambled controls. A consistent decrease in 5HT-2A fluorescence intensity was seen in both cell bodies and in particular neurites following treatment with hu-HTR2A shRNA (Fig. 1D). ImageJ quantification of 5HT-2A immunofluorescence indicated a significant 27% decrease in hu-HTR2A shRNA-AAV9-treated neurons as compared to scrambled-treated (p = .0019) (Fig. 1E). To confirm knockdown of 5HT-2A mRNA, real-time qPCR experiments were undertaken. The results indicated a significant 34% decrease in HTR2A mRNA expression as compared to vehicle controls, p = .012 (Fig. 1F). Relative mRNA levels were not significantly different between vehicle control and scrambled-treated cells (p = .502). Although there was a 24% decrease in mRNA expression between scrambled- and shRNA-treated neurons, this finding did not reach statistical significance (p = .094).

### Adeno-associated virus exposure of human iPSC-differentiated glutamatergic cortical neurons with AAV9-MeCP2-GFP-huHTR2A-shRNA leads to a decrease in spontaneous electrical activity

The serotonin 5HT-2A receptor is the major excitatory receptor subtype in the cortex, therefore, we examined whether exposure of human iPSC-differentiated neurons to AAV9-MeCP2-GFP-huHTR2A-shRNA would lead to a decrease in electrical activity as measured by multi-electrode array (MEA). Multiple parameters were recorded over a 21-day period including the mean burst duration, burst frequency, network bursts, spikes per burst, and synchrony index (see methods for details). As shown in Fig. 2, data are presented over the entire 21-day period (left side) to provide context, and averages of the measures in each of the three treatment conditions over days 12-18 (right side). These three days were selected for repeated measures analysis because by inspection it was evident that following treatment with AAV9-MeCP2-GFP-huHTR2A-shRNA, MEA results before day 12 were essentially flat, and after day 21 the CRO reported that the cells began to lift off the MEA plate and appeared to be dying.

**Fig. 2.**
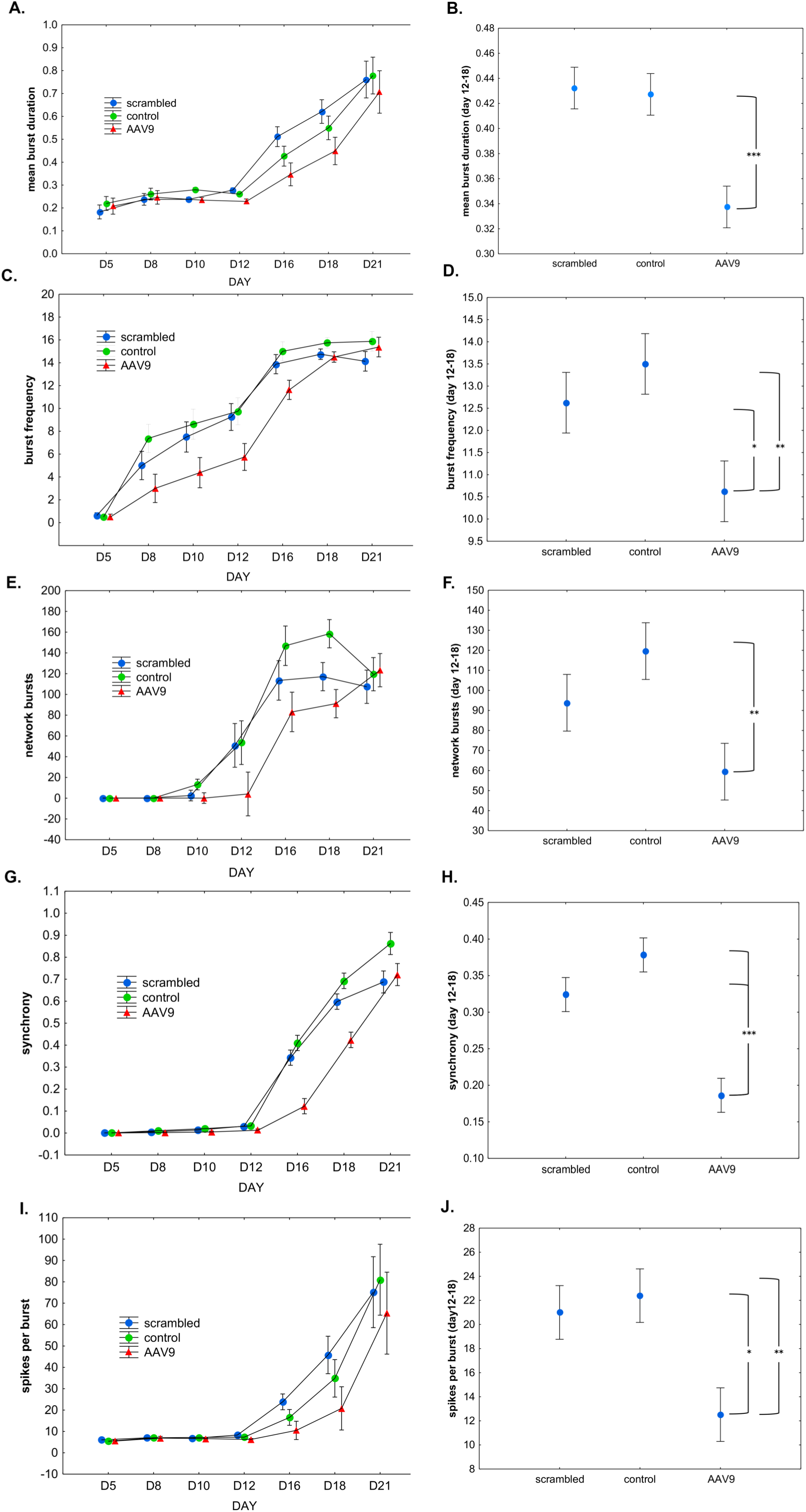
Adeno-associated virus exposure of human iPSC-differentiated glutamatergic cortical neurons with AAV9-MeCP2-GFP-huHTR2A-shRNA leads to a decrease in spontaneous electrical activity. Following plating of neurons, three different conditions were tested starting at Day 5: 1) Vehicle control consisting of PBS only (green circles, labeled “control”); 2) Scrambled AAV9 shRNA at MOI of 3 x 10^5^ (blue circles, labeled “scrambled”); 3) HTR2A hu-shRNA at MOI 3 x 10^5^ (red triangles, labeled “AAV9”). Electrophysiological recordings were acquired three times per week following day 5 *in vitro*. As noted in the text, recording anomalies observed on day 14 were deemed unreliable and were excluded from further analysis. By inspection, it was evident from the AAV9-MeCP2-GFP-huHTR2A-shRNA results that the largest differences noted between the three conditions were on days 12, 16, and 18, and especially so for the synchrony index. Thus, data on these three days were examined for each MEA metric using a repeated measures ANOVA to determine the main effect of “condition,” and then a post-hoc comparison was performed comparing the three conditions, with p-values adjusted via the Fisher LSD method **(A-B)**: Significant differences occurred in mean burst duration between the AAV9 and both of the other two conditions (p< .001). **(C-D)**: Significant differences in average burst frequency were observed between AAV9 vs. the control (p < .01) and scrambled conditions (p < .05). **(E-F)**: A significant difference in average network bursts occurred between AAV9 and the control condition (p < .01) (**G-H**): Significant differences in the synchrony index were observed between both conditions (p< 0.001). (**I-J**): Significant differences in the number of spikes per burst were observed between AAV9 vs. the control (p < .01) and scrambled conditions (p < .05). All data are expressed as the mean ± S.E. Asterisks denote: *p < .05; **p < .01; ***p < .001.

For the 3-day repeated measures analysis the result of interest was whether the average of the three conditions differed from each other. Thus, the factor of “condition” was evaluated, and because there was no *a priori* reason to select those three days, post-hoc comparisons were performed, and the p-values adjusted appropriately using the Fisher LSD (least significant difference) method.

To summarize the data presented in Fig. 2, we observed a decrease in the spontaneous activity of neurons treated with AAV9-MeCP2-GFP-huHTR2A-shRNA in every metric as compared to control conditions and scrambled conditions, and significantly so in every case when compared to control conditions.

### Intranasal adeno-associated virus delivery of mouse AAV9-MeCP2-GFP-mHTR2A-shRNA decreases 5HT-2A receptor expression in vivo

To deliver AAV9-shRNA cargo to mice *in vivo*, we used the nasal route (PCT/US2022/050947). This route bypasses the blood-brain barrier (BBB) and is a practical, non-invasive method. We have previously shown that this route is effective for delivering AAV9 vectors containing CRISPR/Cas9 DNA plasmids ^6^. On day 1, we administered AAV9-MeCP2-GFP-mHTR2A-shRNA intranasally (final concentration ∼1-2.0 x 10^11^ viral particles). Treated mice were then behaviorally assessed 5 and 8 weeks later. After the behavioral test on week 8, mice were sacrificed, and brains were fixed in 4% formalin for immunofluorescence studies or frozen for PCR analyses. The concentration of AAV9 vectors and the time points chosen were based on our previous study ^6^. Compared to vehicle-treated control mice (Fig. 3A-C), widespread neuronal expression of GFP that served as a proxy for mHTRT2A-shRNA expression was evident in subcortical areas including the interpeduncular nucleus (Fig. 3D-F) an area implicated as a major connectome for stress-mediated pathways ^19^ and throughout the olfactory bulb (Fig. 3G-I). As predicted, non-neuronal cells were negative for GFP staining due to the expression of GFPs under MecP2, a specific neuronal promoter. The expression of GFP appeared to localize primarily within the soma of neurons. The strong expression of GFP corresponded with a concomitant decrease in 5HT-2A receptor fluorescence intensity (Fig. 3E), although some staining within apical dendrites was still evident (Fig. 3H). This pattern of staining aligns with our previous results following intranasal treatment of mice with AAV9-CRISPR/Cas9 targeting the HTR2A gene ^6^. ImageJ quantification of 5HT-2A receptor immunofluorescence confirmed a significant decrease in expression (p = .0017) (Fig. 3J).

**Fig. 3.**
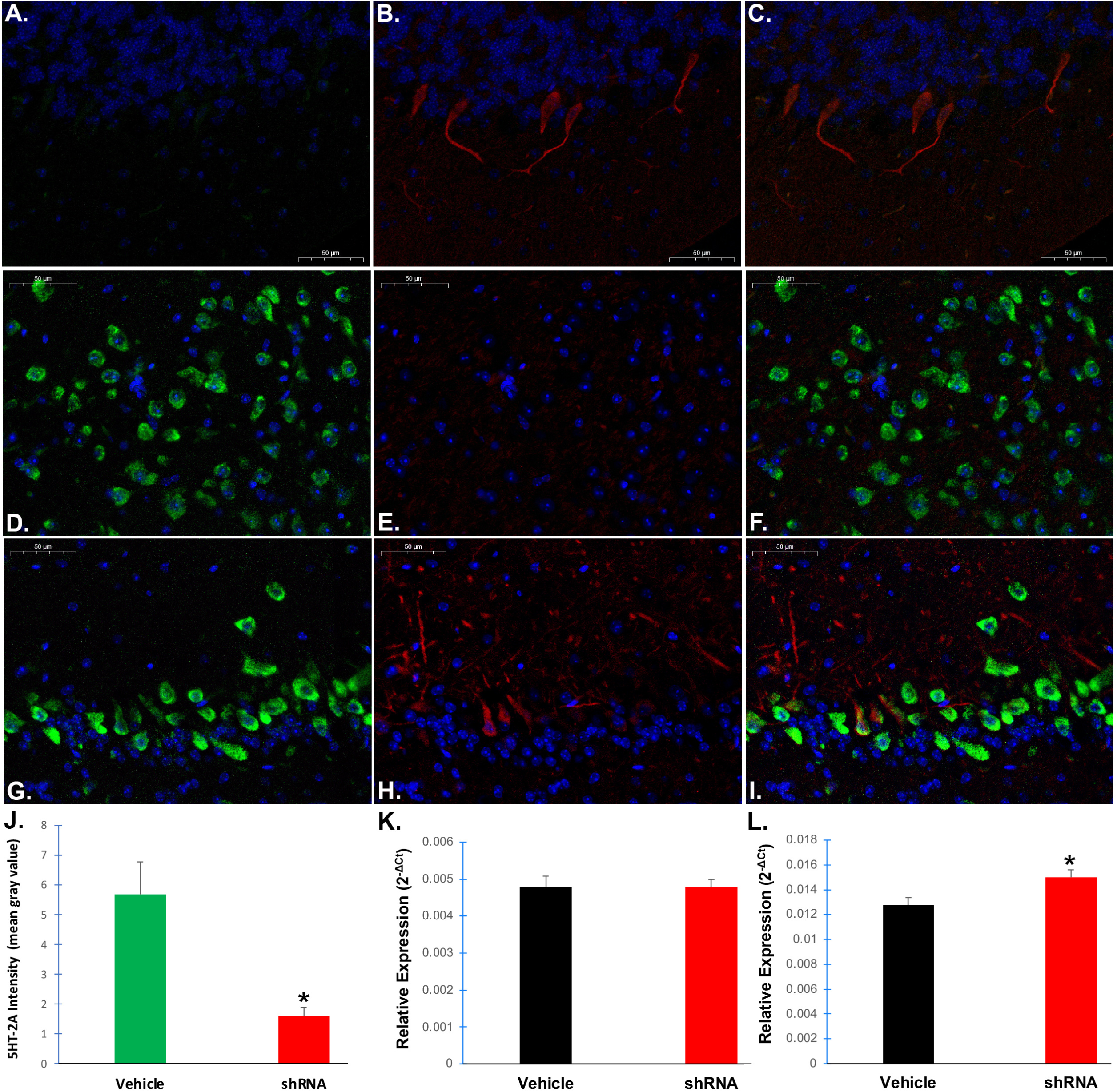
Intranasal adeno-associated virus delivery of AAV9-MeCP2-GFP-mouse HTR2A-shRNA leads to a decrease in 5HT-2A receptor protein expression. Mice were treated with vehicle or AAV vectors on day 1, and 8 weeks later sacrificed, fixed, and 4 µm paraffin-embedded sagittal tissue sections were stained with anti-GFP (green, 1:1,000) or an anti-mouse 5HT2A receptor antibody (red, 1:500). Whole slide imaging was performed using a Pannoramic Midi II scanner (see methods for details). Representative 40X immunofluorescence images from vehicle controls, (cerebellum, **A-C**) or AAV-treated mice are shown (interpeduncular nucleus, **D-F**) and (olfactory bulb, **G-I**). DAPI nuclear stain staining is indicated by blue. Treatment with MeCP2-GFP-mouse HTR2A-shRNA led to a general pattern of less robust staining profile of the 5HT2A receptor in cell body regions and apical dendrites (merged images Panels F and I). Panels A, D, and G represent GFP channel only (a proxy for shRNA expression); Panels B, E, and F represent 5HT-2A receptor protein represent red channel only; Panels C, F, and I represent the merged images. All scale bars represent 50 µm. **(J)**: Quantitative analysis using ImageJ software indicated a significant decrease in 5HT-2A receptor fluorescence intensity of AAV-treated mice (red bar) versus vehicle-controls (green bar). Data represent the mean gray value ±SEM of 2X sagittal brain sections (N=3 for each group). *Denotes significant difference, p-value = .0017. **(K and L)**: Data show the results of qPCR real-time assays to analyze mRNA levels of Htr2a following extraction of total brain RNA from frozen brain (K) or olfactory tissue (L) in either vehicle-controls (black bar) or shRNA treated (red bar). Results display the *relative change* in expression after 8-weeks of treatment with AAV9-MeCP2-GFP-mouse HTR2A. Real-time PCR results represent a total of N=5 animals for each group performed in triplicate ±SEM. *Denotes significant difference between the two groups, p = 0.006 in olfactory mRNA.

Next, we sought to confirm whether intranasal delivery of AAV9 vectors could lead to a decrease in the relative mRNA expression of the *HT2RA* gene. Following treatment of mice on day 1, mice were sacrificed 8 weeks post intranasal treatment and brains and olfactory bulbs were snap-frozen. Total brain or olfactory bulb RNA was extracted from vehicle controls (N=5) or AAV-treated mice (N=5), and real-time PCR was performed as described in the Methods section. There was no significant difference in the relative expression of brain htr2a mRNA between the two groups (p > 0.05) (Fig. 3K), but treatment did lead to a significant difference in the relative expression within the olfactory bulb between the two groups (p = 0.006) (Fig. 3L). Normalized data indicated a 1.2-fold decrease in *htr2a* expression. Although we were unable to demonstrate a robust decrease in htr2a mRNA expression, it is not surprising given the potentially large dilution effect of brain mRNA. Neurons make up less than 10% of the total number of cells in the brain and only a fraction of those neurons would have been infected by AAV9-mHTR2A-shRNA. Moreover, the expression of shRNA from DNA plasmids could attenuate after 8-weeks because the shRNA is maintained as an episomal transgene and not integrated into the host genome ^20^. Indeed, a recent study in mice indicates that vector DNA rapidly decreases 10-fold within neurons over the first 3 weeks following stereotaxic injections of AAV9 vectors into the striatum ^21^.

### Intranasal adeno-associated virus delivery of mouse AAV9-MeCP2-GFP-mHTR2A-shRNA decreases anxiety

Previous studies have shown that serotonin plays a role in pathways associated with stress and anxiety. Therapeutics that reduce serotonin uptake or block serotonin receptors are used to treat anxiety disorders ^22–27^. One serotonin receptor that is thought to be particularly important for anxiety is the 5HT-2A receptor, which is upregulated by stress and mice that lack the 5HT-2A receptor show reduced anxiety ^28^. We tested whether delivering mHTR2A-shRNA to mice could decrease anxiety utilizing the light-dark box test ^29, 30^.

The light/dark box test is a well characterized test used to evaluate the relative anxiety status of mice ^29^. The light/dark paradigm in mice is based on a conflict between the innate aversion to brightly illuminated areas and the spontaneous exploratory activity. If given a choice, mice prefer the dark, and anxiolytic compounds have been found to increase the total duration of time spent in the lit area as well as the number of entries into the lit box ^30^. At 5 weeks post-treatment, AAV9-shRNA-treated mice led to a significant increase compared to vehicle-controls in the time spent in the lit box (34% increase, p < .001) as well as the number of entries into the lit box (22% increase, p = .004) (Fig. 4A and B). It is noteworthy that these results are similar to our previously reported findings using AAV9-CRISPR/Cas9 where we found a 36% increase in time spent in the lit area and a 27% increase in number of entries ^6^. At 8 weeks post-treatment, the effects appear to be slightly attenuated and significance was only observed in the time spent in the lit box (Fig. 4C) but not in the number of entries (Fig. 4D). There was no significant difference in body weight between the two groups, although as in our previous study utilizing AAV9-CRISPR/Cas9, we did observe a trend for a slight decrease in weight in the AAV9-shRNA-treated mice (Fig. 4E).

**Fig. 4.**
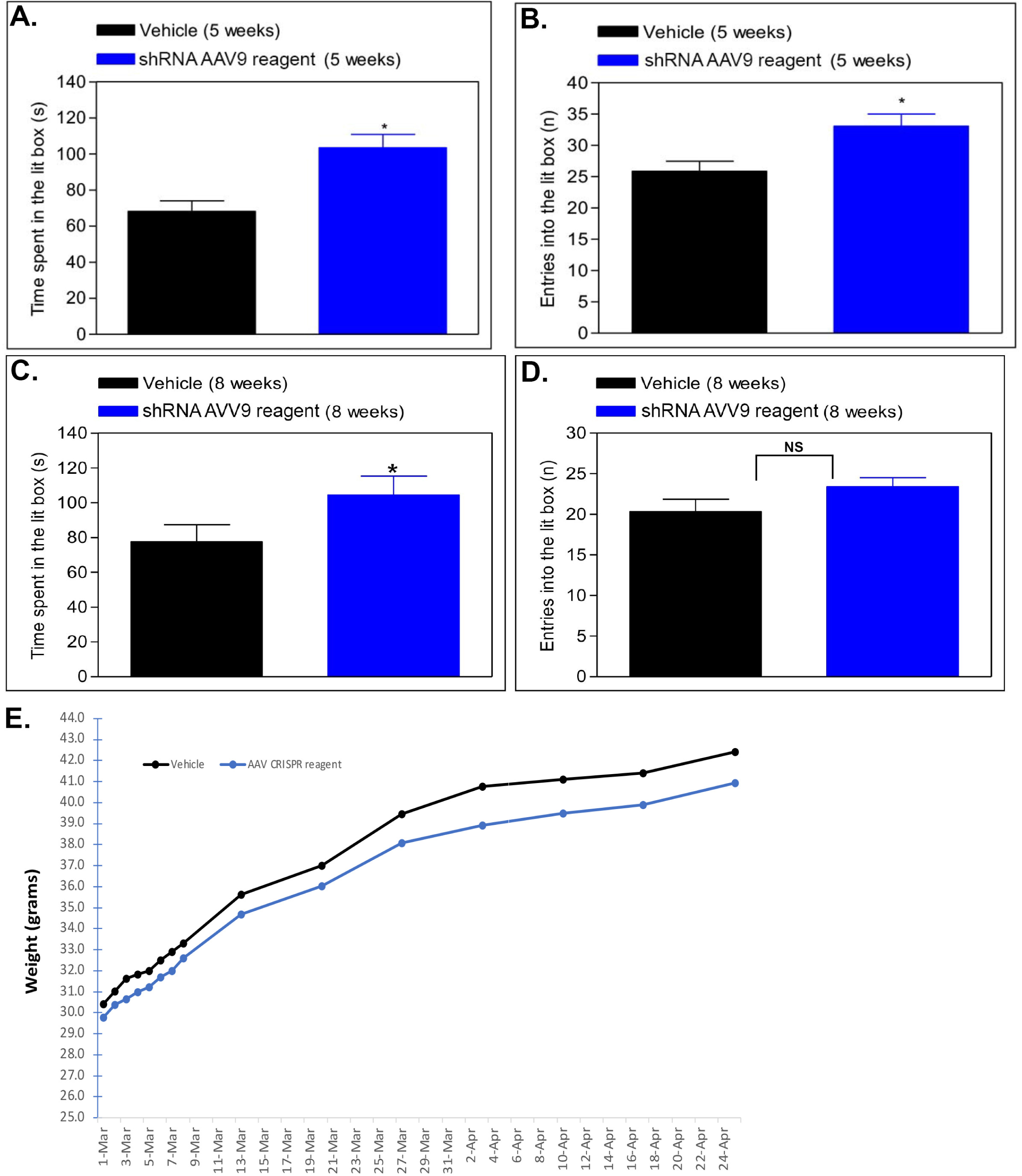
Intranasal adeno-associated virus delivery of AAV9-MeCP2-GFP-mouse HTR2A-shRNA decreases anxiety. Mice were treated intranasally on day 1 with 2.0 x 10^11^ viral particles and compared to vehicle-controls either at 5-weeks (**A and B**) or 8-weeks (**C and D**) later in the light dark test to evaluate the relative anxiety status of mice. Results for 5-week AAV9-shRNA-treated mice indicated a 34% increase in the time spent in the lit box (N=15, p-value <.001) as well as an 22% increase in the number of entries into the lit box (p = .004). Following a retesting of the same animals at 8-weeks, a slightly diminished response in both parameters was noted, however there was still a significant increase in the time spent in the lit box **(C)** (p-value = 0.04). The number of entries into the light box at 8-weeks just missed statistical significance **(D)** (p-value = .058). **(E)**: Although there was a trend for slightly lower weight in AAV9-shRNA-treated mice, data indicated no significance between the two groups (p = 0.49). *Denotes statistical significance between the two groups (p-value <0.05). NS denotes non-significant (p-value>0.05).

### Intranasal delivery of either AAV9-CRISPR/Cas9 or AAV9-shRNA improves memory

Whether blocking or downregulating the 5HT-2A receptor is expected to improve memory outcomes is not well established. To address this, we employed two different animal model species using two different memory tests. In the first test, we treated aged mice with our AAV9-CRISPR/Cas9 construct to decrease 5HT-2A receptor density through selective knockout of the *HTR2A* gene ^6^. Mice were treated on day 1 and assessed using the T-maze continuous alternation task 5 weeks later (see Methods for details).

As shown in Fig. 5, compared to the vehicle-control group, AAV9-CRISPR/Cas9-treated mice showed a highly significant increase in the number of spontaneous alterations (p = 0.0007). This finding equates to a 104% increase in memory. Examination of CA2 and CA3 regions of the hippocampus an area of the brain implicated in memory indicated 5HT-2A receptor staining within neuronal processes but not cell bodies of vehicle-controls (Fig. 5B). In contrast, for treated animals, 5HT-2A staining was abolished with a corresponding strong expression of GFP (a proxy for guide RNA expression) within the CA2 and CA3 regions of the hippocampus (Fig. 5C). Taken together, these data suggest that specific knockdown of the 5HT-2A receptor using CRISPR has the potential to be a promising therapeutic approach for the treatment of age-related cognitive decline.

**Fig. 5.**
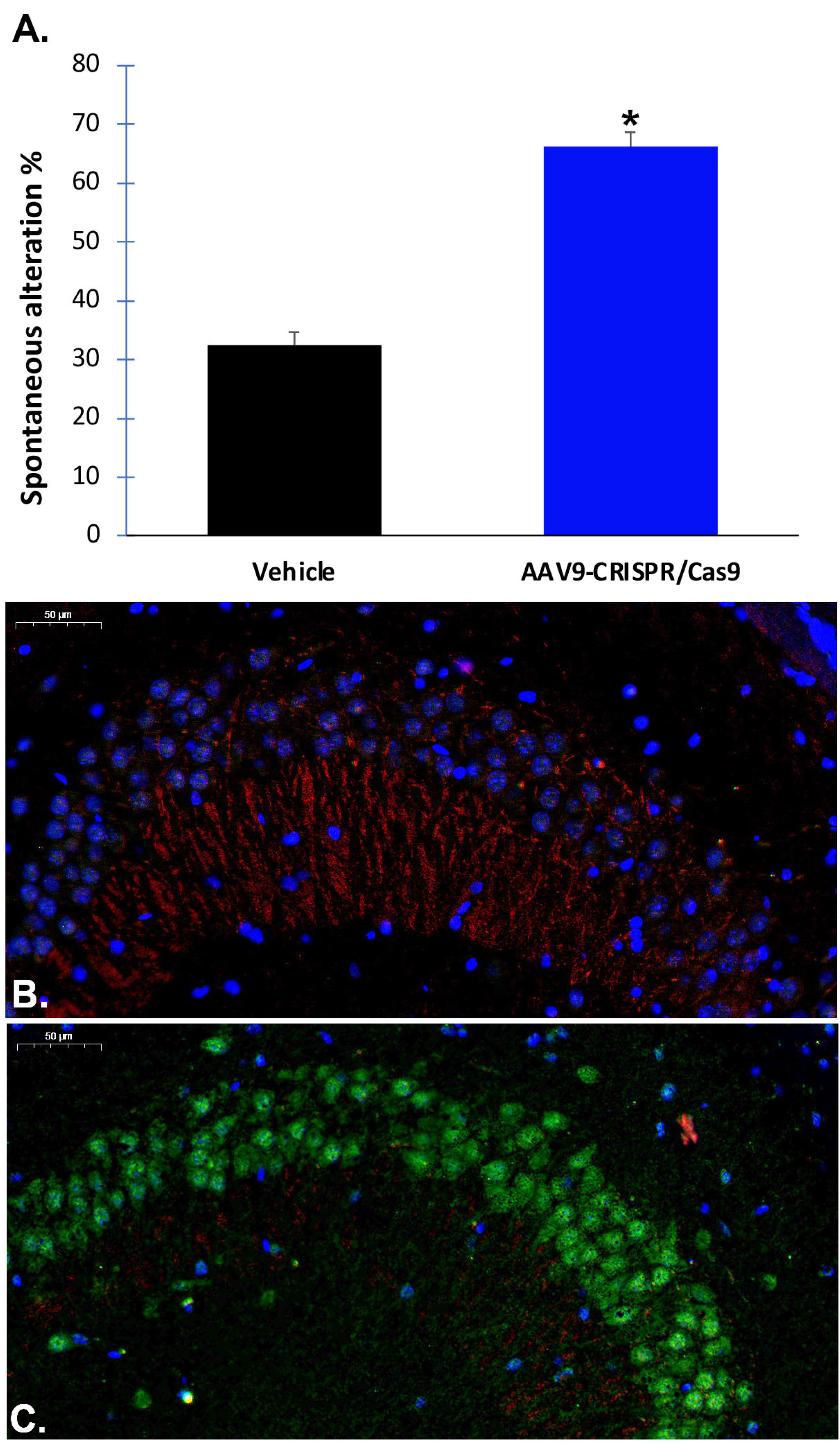
Intranasal adeno-associated virus delivery of co-packaged *Cas9* DNA and a *HTR2A*-targeting guide RNA improves memory in aged mice. **(A)**: Aged 12-month-old mice were treated intranasally with vehicle or with 2.0 x 10^11^ viral particles on day 1, and 5-weeks later tested behaviorally using a spontaneously alteration memory test. Mice treated with AAV9-CRISPR/Cas9 showed a significant increase in the percent spontaneous alterations (p-value = .0007, N=15 mice per group, asterisk, blue bar). **(B)**: Representative, merged immunofluorescence image of vehicle-control animals depicting the presence of 5HT-2A receptor protein labeling in apical dendrites in the CA2/CA3 region of the hippocampus. The blue staining reflects nuclear staining with DAPI. As expected, there was no expression of GFP in vehicle controls. **(C)**: Identical to Panel B with the exception that the merged image is from a AAV9-CRISPR/Cas9-treated mouse brain. In this case, strong GFP labeling was observed in cell bodies while there was an observed decrease in 5HT-2A receptor fluorescence. Images are representative of 3 separate mice for each group.

To confirm these results, we tested a different species, rats, using a novel object recognition test following treatment with AAV9-mHTR2A-shRNA. In this case, 2-month-old rats were treated with AAV9-mHTR2A-shRNA on day one and tested 3- and 5-weeks later. The novel object recognition task is used to assess short-term memory, intermediate-term memory, and long-term memory in rats ^16^. The task is based on the natural tendency of rats to preferentially explore a novel versus a familiar object, which requires memory of the familiar object (see Methods for details). As shown in Fig. 6, at 3-weeks significant increases in both the contact-recognition index (92% increase, p < 0.000003) and time-recognition index (73%, p< 0.0003) were observed (Fig. 6C and D). Importantly, there were no differences in the amount of time spent during the learning phase (p > 0.05) (Fig. 6A and B), therefore, these results represent a true increase in memory retention. In addition, there was no significant difference in weight at 3-weeks (p = 0.463). A Bayesian analysis for memory data at 3 weeks indicated a Bayes Factor in favor of the hypothesis that the AAV9-HTR2A-shRNA treatment would produce better memory results than the control treatment for the novel object contacts and time metrics was 4,377 and 106, respectively. Considering that the rule of thumb for a Bayes Factor of 100 is considered “decisive evidence,” we conclude that the treated rats had superior memory as compared to the vehicle-controls.

**Fig. 6.**
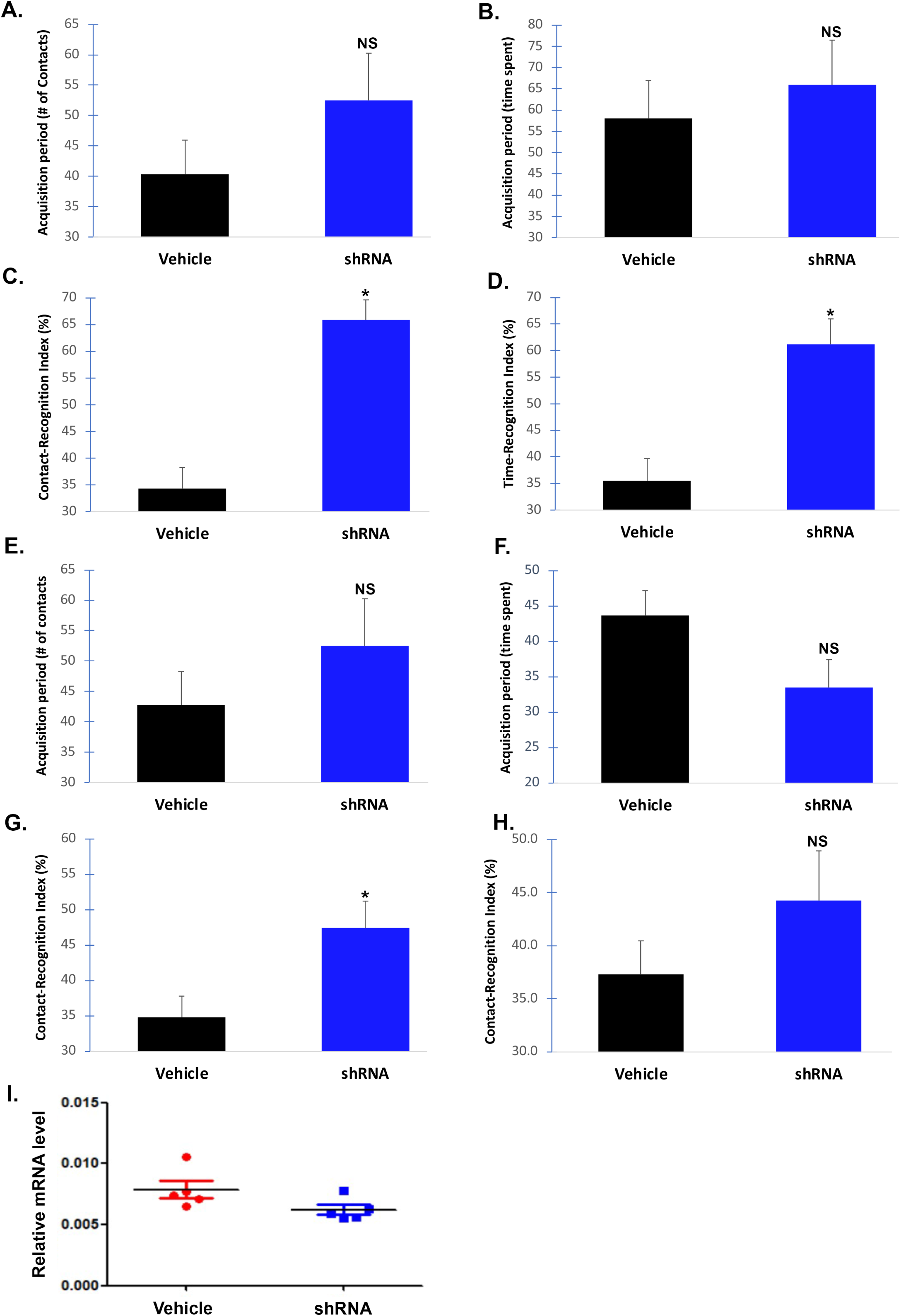
Intranasal adeno-associated virus delivery of AAV9-MeCP2-GFP-mouse HTR2A-shRNA improves memory in rats. The target sequence used to synthesize the shRNA is 100% conserved between mice and rats. To test whether shRNA-knockdown of the rat 5HT-2A receptor improves memory, Wistar rats (12 animals per group) were randomly assigned to two different groups consisting of vehicle- or AAV9-MeCP2-GFP-mHTR2A-shRNA. Following treatment on day 1, animals were assessed behaviorally 3- (**A-D**) and 5-weeks later (**E-H**). Details of the novel object recognition test can be found in the methods. At 3 weeks, there was no significance difference in the time spent or the number of physical contacts with the novel object during the acquisition (training) period (p > 0.05) (**A and B**). Rats were tested 24-hours later, and AAV9-treated rats (blue bars) showed a significant increase in both the contact-recognition index (p < 0.000003) (C) and the time recognition-index (p <0.0003) (D). The same groups of rats were retested at 5-weeks and again no significant difference was noted in the time spent or the number of physical contacts with the novel object during the acquisition (training) period (p > 0.05) (**E and F**). Twenty-four hours later there was a significant increase in the contact-recognition index (p < 0.01) (**G**), however, there was no significant difference in the contact-recognition index (p = 0.114) (**H**). (**I**): Data show the results of qPCR real-time assays to analyze mRNA levels of Htr2a following extraction of rat brain RNA. Results display relative mRNA levels after 5-weeks post-treatment with AAV9-HTR2A-shRNA. Real-time PCR results represent a total of N=5 animals for each group performed in triplicate ±SEM. *Denotes significant difference between the two groups, p = 0.038. (**J**): Representative, merged immunofluorescence image of vehicle-control animals depicting the presence of 5HT-2A receptor protein labeling (red fluorescence) within the olfactory bulb. The blue staining reflects nuclear staining with DAPI. As expected, there was no expression of GFP in vehicle controls (J, left panel) while strong GFP labeling was observed in cell bodies of neurons of shRNA-treated rats (J, right panel). Images are representative of 3 separate mice for each group.

At 5-weeks post treatment the enhanced memory effects were attenuated and only contact-recognition index showed a significant 36% increase (p = 0.008) (Fig. 6G and H). The contact-recognition index was 19% higher than vehicle-control rats but did not reach statistical significance (p = 0.114). To test whether treatment led to a decrease in 5HT-2A receptor mRNA levels, qPCR was performed. A significant decrease in relative mRNA levels was observed 5-weeks post treatment (p = .038) (Fig. 6I). We also were able to confirm the expression of the HTR2A-shRNA within the olfactory bulb using GFP fluorescence as a proxy (Fig. 6J).

Considered together, for the novel object recognition test the results showed a highly significant improvement in memory retention at week 3 and a significant improvement at 5 weeks. The attenuation of memory retention at 5-weeks may be a result of an increase in degradation of the plasmid DNA, an extinction in memory because of the same rats being tested twice in a span of 2 weeks, or a combination of the two.

## Discussion

According to the American Psychiatric Association, anxiety disorders make up the most common type of mental disorders, affecting nearly 30 percent of adults at some point in their lives. Chronic anxiety in turn can impact memory and as a result, persistent anxiety and memory impairments are inextricably linked. Indeed, anxiety disorders are interrelated and inseparable with memory loss, and anxiety is likely an early predictor of future cognitive impairment ^31–35^. Numerous studies now support that anxiety and depression are key co-morbidities associated with AD ^36–39^. In the present study, we demonstrate that short-hairpin RNAs (shRNA) targeting knockdown of the human *HTR2A* gene can modify neuronal circuits underlying anxiety and memory. *In vitro*, using human iPSC-differentiated neurons, shRNA targeting the HTR2A RNA transcript led to decreased expression of the 5HT-2A mRNA and receptor protein as well as a concomitant decrease in spontaneous electrical activity of cortical neural networks. *In vivo*, application of AAV9-shRNA by intranasal delivery led to a decrease in 5HT-2A receptor density and a significant decrease in anxiety following 5-weeks post-treatment in mice. However, this anxiolytic effect was attenuated at 8-weeks post-treatment. There are several possible explanations for the decreased efficacy of AAV9-shRNA at 8 weeks of treatment. First, it is possible that the plasmid DNA within infected cells was degraded, which is supported by the lack of difference in mRNA levels between the two groups at 8 weeks (Fig. 3K). Second, the diminished response could be due to habituation, because the same mice were used for both the 5- and 8-week light/dark tests. Finally, it is possible that both factors are at play simultaneously. In summary, we found that intranasal delivery of AAV9-htr2a-shRNA, which reduces the expression of the 5HT-2A receptor, significantly decreased anxiety of mice in the light/dark box test.

Studies employing specific 5HT-2A receptor antagonists, including M200907, ritanserin, ketanserin, TCB-2, and risperidone, give mixed results with some increasing acquisition or consolidation while others do not (for review see ^40^). On the other hand, studies examining the knocking down of the 5HT-2A receptor have shown improvement in memory in rodent models ^9–11^. In humans, administration of mianserin (15 mg/day), an agent with marked 5HT-2A antagonism, improved memory, learning and attention ^41^. These conflicting results may reflect the different underlying neural mechanisms involved in different types of memory as well as the dose of the administered 5HT-2A antagonist. Because of the uncertainty of the role of 5HT-2A receptor in memory, in the current study we tested whether downregulation of the 5HT-2A receptor, either through CRISPR/Cas9 or shRNA, could impact memory. Our *in vivo* experiments showed that knockdown of the 5HT-2A receptor led to a significant improvement in memory in both an aged-mouse model as well as in 2-month-old rats. How does a decrease in 5HT-2A receptor density contribute to a significant increase in memory? Several possibilities exist, including a decrease in anxiety, which may itself promote better memory outcomes. Knockdown of the 5HT-2A receptor within the hippocampus may be another proposed mechanism for these findings. Indeed, we found a general pattern of guide RNA expression within the CA2/CA3 region of the hippocampus of CRISPR/Cas9-treated mice and an apparent decrease expression of the 5HT-2A receptor in the same region, particularly within apical dendrites of glutamatergic neurons. Previous studies have demonstrated 5HT-2A receptor mRNA expression in the CA3 region of the hippocampus ^42,43^. Because the 5HT-2A receptor is excitatory, downregulation of this receptor in apical dendrites within the hippocampus may improve memory through the modulation of hippocampal neuronal and glial oscillatory rhythm ^44, 45^. Indeed, reduction of hippocampal hyperactivity has been shown to improve cognition in amnestic mild cognitive impairment ^46^. While molecular mechanisms of memory formation, retention and recalling in hippocampal neurons are not yet fully understood, recent evidence shows that individual neurons code discrete memories using either a rate code or a temporal firing code ^47^. Presently, however, our knowledge of the complex interplay of serotonin modulation of hippocampal glutamatergic neurons is at a very early stage.

In conclusion, our study highlights the immense potential of short-hairpin RNAs (shRNA) as a precision-based therapeutic approach for neurodegenerative disorders. While the use of shRNA molecules to modify neuronal circuits underlying specific behaviors is a relatively unexplored territory, our findings shed light on their promising efficacy. Through the design of shRNA targeting the human HTR2A gene, we demonstrated significant outcomes both *in vitro* and *in vivo*. *In vitro* experiments using iPSC-differentiated neurons revealed that knockdown of the 5HT-2A receptor led to a notable decrease in spontaneous electrical activity, emphasizing the pivotal role of this receptor in neural network dynamics.

*In vivo*, intranasal delivery of AAV9 vectors carrying shRNA resulted in a remarkable reduction in anxiety-like behavior in mice, addressing a critical aspect of mental health. Additionally, our treatment led to a substantial improvement in memory in both mice and rats, with memory enhancements reaching up to 104% and 92%, respectively, compared to vehicle-treated animals. These results suggest that targeting the *HTR2A* gene can simultaneously alleviate chronic anxiety and ameliorate age-related cognitive decline.

Perhaps equally significant is the development of a non-invasive shRNA delivery platform capable of bypassing the blood-brain barrier. This achievement opens up a wide range of possibilities for the treatment of various neurological and mental disorders. In particular, the novel therapeutic approach we have demonstrated holds great promise for addressing the intertwined issues of chronic anxiety and cognitive decline, offering hope for improving the quality of life for individuals affected by these conditions.

