## Supplemental Material for "Intranasal Delivery of shRNA to Knockdown the 5HT-2A receptor Enhances Memory and Alleviates Anxiety"

### Supplementary Information

A.

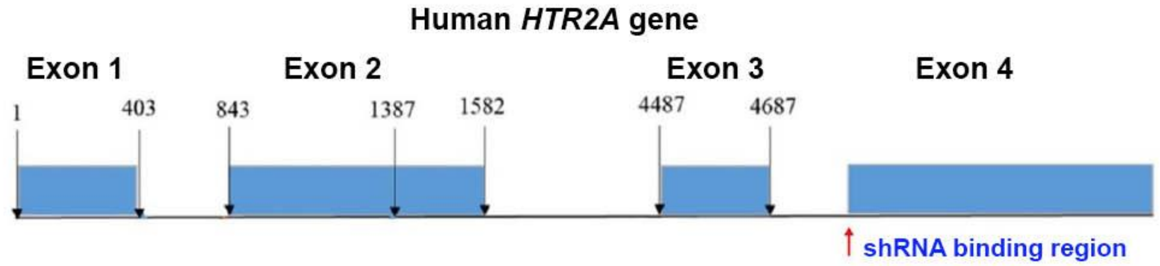


B.

| shRNAmir Vector | KD % | shRNAmir sequence |
|---|---|---|
| Empty Vector control | 0% | |
| shmir#1 | 76% | GCT GAGCACGTCCAGGTAAATCCAGGTTTTGG CCAC TGACTGACCTGGATTTCTGGACGTGCT CAG |
| shmir#2 | 85% | GCT GTACTGATATGGTCCAAACAGCGTTTTGG CCAC TGACTGACGCTGTTTGCCATATCAGTA CAG |
| Shmir#3 ** | 87% | GCT GTTCTGAAGACAAAGAACTCTGGTTTTGG CCAC TGACTGACCAGAGTTCTGTCTTCAGAA CAG |
| shmir#4 | 72% | GCT GCAAACAAACACATTGAGCAGGGTTTTGG CCAC TGACTGACCCTGCTCAGTGTTTGTTTG CAG |

C.

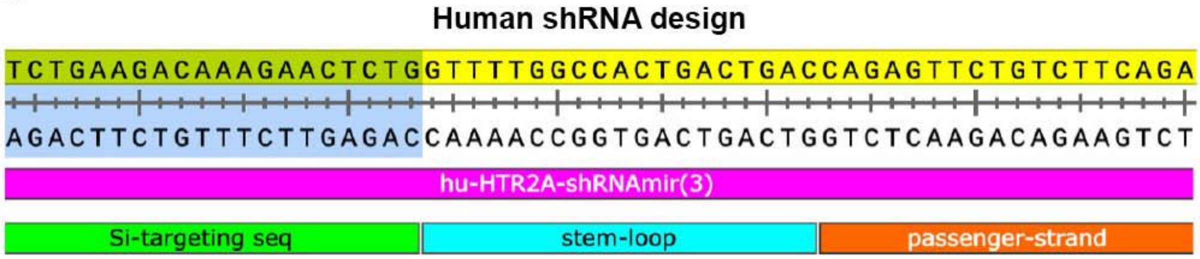


D.

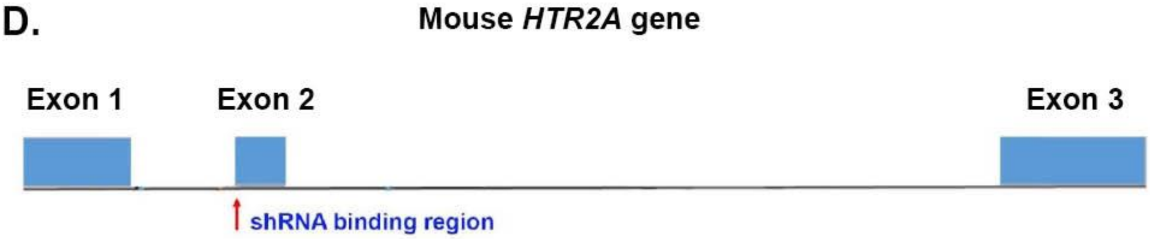


E.

| shRNAmir Vector | KD % | shRNAmir sequence |
|---|---|---|
| Empty Vector control | 0% | |
| Scrambled shRNAmir control | 0% | |
| shmir#1 | 68% | GCT GTCAATTGTCCAGTTCGAGGCTGTTTTGG CCAC TGACTGACAGCCTCGATGGACAATTGA CAG |
| shmir#2 | 61% | GCT GTGGAGATGAAGAATGGAGAGGGTTTTGG CCAC TGACTGACCCTCTCCACTTCATCTCCA CAG |
| Shmir#3 | 62% | GCT GAAACCCAGCAGCATATCAGCTGTTTTGG CCAC TGACTGACAGCTGATACTGCTGGGTTT CAG |
| Shmir#4 ** | 77% | GCT GAGCACATCCAGGTAAATCCAGGTTTTGG CCAC TGACTGACCTGGATTTCTGGATGTGCT CAG |

**Supplementary Fig. 1. Targeting strategy to knock down the human and mouse *HTR2A* gene using shRNA. (A)**: The *HTR2A* human gene located on chromosome 13q14-21 spans 66 kilobases and consists of 4 exons. At least 2 major isoforms are generated from alternative splicing between exons 1-2. Therefore, we chose to construct a shRNA sequence targeting the beginning of exon 4, which would lead to degradation of the largest coding exon and potentially downregulation of the two major isoforms of the human *HTR2A* gene. **(B)**: Four different shRNA constructs were validated and screened *in vitro*. Shmir#3, which produced an 87% knockdown was selected to be used for viral production.

**(C)**: The design of the human shRNA consists of three parts: 1) The target sequence shown (shaded sequence) and depicted in green; 2) The stem-loop region depicted in light blue; 3) The passenger strand sequence shown in orange. Details on the construction of this shRNA can be found in the methods section. **(D)**: The *HTR2A* mouse gene encodes a single protein-coding transcript, *Htr2a-201*, located on chromosome 14. A similar strategy as to constructing the human shRNA was employed to design the mouse version. In this case, the target sequence was designed to bind at the beginning of exon 2. Silencing of exon 2 would prevent the 5HT-2A receptor protein from being produced. **(E)**: Four different shRNAs were tested for potential knockdown *in vitro*. Shmir#4 was chosen for viral production following demonstration of a 77% knockdown with no knockdown observed either with the empty vector control or a scrambled shRNA control.

Table 1: Seeding medium for culturing human iPSC differentiated neurons

| | | Component | Stock Conc. | Final Conc. | 1 Plate Volume | 2 Plate Volume | 5 Plate Volume |
|---|---|---|---|---|---|---|---|
| Seeding Medium | 1 | DMEM/F12 Medium | 1X | 0.5X | 9.5 mL | 19 mL | 47.5 mL |
| | 2 | Neurobasal Medium | 1X | 0.5X | 9.5 mL | 19 mL | 47.5 mL |
| | 3 | B27 Supplement | 50X | 1X | 400 µL | 800 µL | 2 mL |
| | 4 | N2 Supplement | 100X | 1X | 200 µL | 400 µL | 1 mL |
| | 5 | GlutaMAX | 200 mM | 0.5 mM | 50 µL | 100 µL | 250 µL |
| | 6 | BDNF | 10 µg/mL | 10 ng/mL | 20 µL | 40 µL | 100 µL |
| | 7 | GDNF | 10 µg/mL | 10 ng/mL | 20 µL | 40 µL | 100 µL |
| | 8 | TGF-β1 | 1 µg/mL | 1 ng/mL | 20 µL | 40 µL | 100 µL |
| | 9 | Geltrex | 15 mg/mL | 15 µg/mL | 200 µL (of 1:10) | 400 µL (of 1:10) | 1 mL (of 1:10) |
| | 10 | Glut Neuron Seeding Supplement | 1000X | 1X | 20 µL | 40 µL | 100 µL |
| | 11 | Supplement K | 1000X | 0.5X | 10 µL | 20 µL | 50 µL |

Table 2: Day 4 medium for culturing human iPSC differentiated neurons

| | | Component | Stock Conc. | Final Conc. | 1 Plate Volume | 2 Plate Volume | 5 Plate Volume |
|---|---|---|---|---|---|---|---|
| Day 4 | 1 | DMEM/F12 Medium | 1X | 0.25X | 4.8 mL | 9.6 mL | 20 mL |
| | 2 | Neurobasal Medium | 1X | 0.25X | 4.8 mL | 9.6 mL | 20 mL |
| | 3 | BrainPhys Medium | 1X | 0.5X | 9.6 mL | 19.2 mL | 48 mL |
| | 4 | B27 Supplement | 50X | 1X | 400 µL | 800 µL | 2 mL |
| | 5 | N2 Supplement | 100X | 1X | 200 µL | 400 µL | 1 mL |
| | 6 | GlutaMAX | 200 mM | 0.5 mM | 50 µL | 100 µL | 250 µL |
| | 7 | BDNF | 10 µg/mL | 10 ng/mL | 20 µL | 40 µL | 100 µL |
| | 8 | GDNF | 10 µg/mL | 10 ng/mL | 20 µL | 40 µL | 100 µL |
| | 9 | TGF-β1 | 1 µg/mL | 1 ng/mL | 20 µL | 40 µL | 100 µL |
| | 10 | Day 4 Supplement | 1000X | 1X | 20 µL | 40 µL | 100 µL |
| | 11 | Supplement K | 1000X | 0.5X | 10 µL | 20 µL | 50 µL |

Table 3: Maintenance media for culturing human iPSC differentiated neurons

| | | Component | Stock Conc. | Final Conc. | 1 Plate Volume | 2 Plate Volume | 5 Plate Volume |
|---|---|---|---|---|---|---|---|
| Maintenance Medium | 1 | BrainPhys | 1X | 0.5X | 19.3 mL | 38.6 mL | 96.5 mL |
| | 2 | B27 Supplement | 50X | 1X | 400 µL | 800 µL | 2 mL |
| | 3 | N2 Supplement | 100X | 1X | 200 µL | 400 µL | 1 mL |
| | 4 | GlutaMAX | 200 mM | 0.5 mM | 50 µL | 100 µL | 250 µL |
| | 5 | BDNF | 10 µg/mL | 10 ng/mL | 20 µL | 40 µL | 100 µL |
| | 6 | GDNF | 10 µg/mL | 10 ng/mL | 20 µL | 40 µL | 100 µL |
| | 7 | TGF-β1 | 1 µg/mL | 1 ng/mL | 20 µL | 40 µL | 100 µL |
